# Super-resolution microscopy of nanoscopic alpha-synuclein aggregates in brain samples indicates a subset of cells have disrupted protein homeostasis prior to Lewy body formation in Parkinson’s disease

**DOI:** 10.1101/2025.09.16.676632

**Authors:** Emre Fertan, John S. H. Danial, Stephen Neame, Jeff Y. L. Lam, Matthew W. Cotton, Melanie Burke, Zengjie Xia, Yunzhao Wu, Ben Powney, Yoichi Imaizumi, Annelies Quaegebeur, Georg Meisl, James Staddon, David Klenerman

## Abstract

Nanoscopic aggregates of alpha-synuclein have been observed in Parkinson’s disease (PD). However, the processes that occur *in-vivo* leading to the formation of these small aggregates are not well understood. We used ultra-sensitive single-molecule methods including SIMOA and super-resolution microscopy to quantify and characterise alpha-synuclein aggregates harvested from human brain samples alongside the Line 61 mouse model using different tissue processing methods. While aggregate numbers did not differ between PD and control samples, larger aggregates were detected in PD brain samples. Moreover, different sub-populations of aggregates were obtained by different extraction methods, with diffusible and membrane-bound aggregates producing a more pronounced difference between disease and control samples. Our data suggests that alpha-synuclein aggregates slowly in the brain, leading to formation of larger aggregates in a sub-set of cells.

## Introduction

Affecting 4% of the population over the age of 80, Parkinson’s disease (PD) is the most common neurodegenerative disorder leading to movement deficits(1). With motor symptoms of rigidity, bradykinesia, and resting tremor(2), accompanied by speech dysfunction(3) - and in some cases cognitive impairment(4), PD significantly decreases the quality of life(5) and shortens the lifespan(6). While the exact cause(s) and neuropathological mechanisms of PD are still being studied, the loss of dopaminergic neurons in the substantia nigra pars compacta(7) as well as the aggregation and accumulation of alpha-synuclein (□Syn)(8,9) have been well documented in PD brains.

As a synaptic protein, □Syn is required for neurotransmitter release by aiding the formation of the soluble N-ethylmaleimide-sensitive factor attachment protein receptor (SNARE) complex(10). However, in PD, □Syn forms large, insoluble aggregates called Lewy bodies(8), concentrating in regions where neuronal loss occurs, such as the substantia nigra(11). While the microscopic Lewy bodies consisting of fibrillar □Syn are the most commonly studied forms of □Syn aggregation and their atomic structures have been resolved using cryogenic electron microscopy (cryoEM); their role in the pathogenesis PD has not been fully established. Meanwhile, some recent studies have shown that the earlier forms of aggregation, namely the nanoscopic aggregates - which are highly heterogeneous in size and shape(12), may be more toxic than the Lewy bodies(13). Sometimes termed “oligomeric □Syn”, these smaller aggregates inhibit neurotransmitter vesicle docking and release(14,15). Our group has shown that these diffusible □Syn aggregates are present in the amygdala of the brains of PD patients and notably also the brains from unaffected individuals, but the diffusible aggregates differ in size between PD and control cases, and the aggregates under 200 nm in length are neurotoxic(16).

While various studies have demonstrated the toxic properties of small aSyn aggregates, the exact species of these nanoscopic aggregates found in PD cases remains elusive as it is challenging to characterise and study these aggregates. Firstly, the small aSyn aggregates are highly heterogenous, low in abundance, and below the diffraction-limit of light, making them impossible to be studied and characterised with traditional microscopy techniques and bulk analysis methods, as they are smaller and less abundant than the resolution limit and sensitivity of these techniques. Single-molecule and super-resolution detection methods are therefore required to resolve individual aggregates, so they can be quantified and characterised. Moreover, the methods used to process the brain tissue and harvest the aggregates often yield a subset of the aggregates present in the tissue and may alter their properties and size distribution. This can bias observations towards the species which are dominant for a particular extraction method. For instance, Hong *et al*.(17) demonstrated that the method used to process post-mortem Alzheimer’s disease brain samples impacts the type of beta-amyloid aggregates harvested. While gently soaking the samples gathered a small sub-subsection of the aggregates, that are termed “diffusible”(18) these aggregates were highly toxic and capable of inducing long-term potentiation deficits(17,19). Meanwhile, the mechanical disruption of the cellular integrity can harvest the aggregates freely-present in the cytosol yet cannot solubilise the membrane-bound aggregates. Since aSyn is known to interact with membranes and show oligomerisation on lipid bilayers(20–22), detergents such as Triton X-100 (TrX) are needed to harvest the aggregates bound to membranes(23). Lastly, harsher anionic detergents such as N-laurylsarcosine (sarkosyl) are known to de-stabilise fibrillar □Syn aggregates and release monomeric and aggregated species(24).

In order to compare the nanoscopic aggregates harvested by different extraction methods and identify the “disease relevant” species, we have combined single-molecule and super-resolution detection methods with advanced data modelling. We first applied these methods to human post-mortem PD and control brain samples from the orbitofrontal cortex at Braak stages 4 and 5, which is prior to Lewy body pathology in this region. We then studied the commonly-used Line 61 mouse model of PD, to determine if the mouse model mimics the human disease in terms of the disease-relevant small aggregates. Finally, we employed a mathematical model which we have recently developed to interpret aggregate size distributions in terms of the underlying molecular mechanisms(25) to compare the PD and control samples, along with the Line 61 mice.

## Results

The small □Syn aggregates studied here are mostly below the diffraction-limit of light (~200 nm), thus cannot be resolved and characterised using traditional light microscopy techniques such as immunohistochemistry. Moreover, their high heterogeneity and low abundance makes it difficult to separate the disease-relevant species from the larger aggregates and monomers with bulk analysis methods such as ELISA. As such, single-molecule detection methods are required to quantify and morphologically characterise individual aggregates with high precision, and to compare them between different extraction methods. Here, we used an ultra-sensitive single molecule array (SIMOA) assay developed by our group that has been shown to quantify the amount of □Syn aggregates specifically (excluding monomers) in the samples, with femtomolar sensitivity(26). Subsequently, the aggregates were characterised in terms of their size and shape, using a combination of single-molecule pulldown (SiMPull) and DNA-PAINT imaging, previously developed and used by our group to characterise beta-amyloid(27) and tau aggregates(28). In both assays the same monoclonal antibody was used to capture and detect the targets, thus aggregates are studied - excluding the monomeric □Syn in the samples as the single epitope is taken up by the capture antibody, making it impossible for the detection antibody to bind to the monomeric target. This was confirmed experimentally by denaturation experiments and super-resolution imaging for the SIMOA and SiMPull respectively. Moreover, the total number of localisations (imaging strands bound to the aggregates) can be calculated in DNA-PAINT, positively correlating with the total target mass in the sample(29). The results of these experiments are presented below, and the full results of statistical tests (including degrees of freedom, p-values, and CI’s) are provided in the **supplemental results file**.

### 3.1 □Syn aggregate concentration in human PD and control brain

We processed age- and sex-matched PD and control post-mortem human orbito-frontal cortex samples with serial extractions and compared the quantity as well as the morphology of the □Syn aggregates (**see Figure 2 for sample images**) using a purpose-built SIMOA aggregate assay and DNA-PAINT super-resolution microscopy, respectively. Strikingly, the □Syn aggregate quantity did not differ between the PD and the control samples (**Figure 3a**) as the number of nanoscopic aggregates were not increased in the orbitofrontal cortex of the PD brains. Meanwhile, there was a significant difference between aggregate quantities obtained using the different extraction methods. The sarkosyl soluble fraction contained the least number of aggregates for both disease and control samples, followed by the soaked and TrX extracted fractions, and the homogenised samples contained the highest concentration of □Syn aggregates (**Figures 3a & 4**).

**Figure 1.**
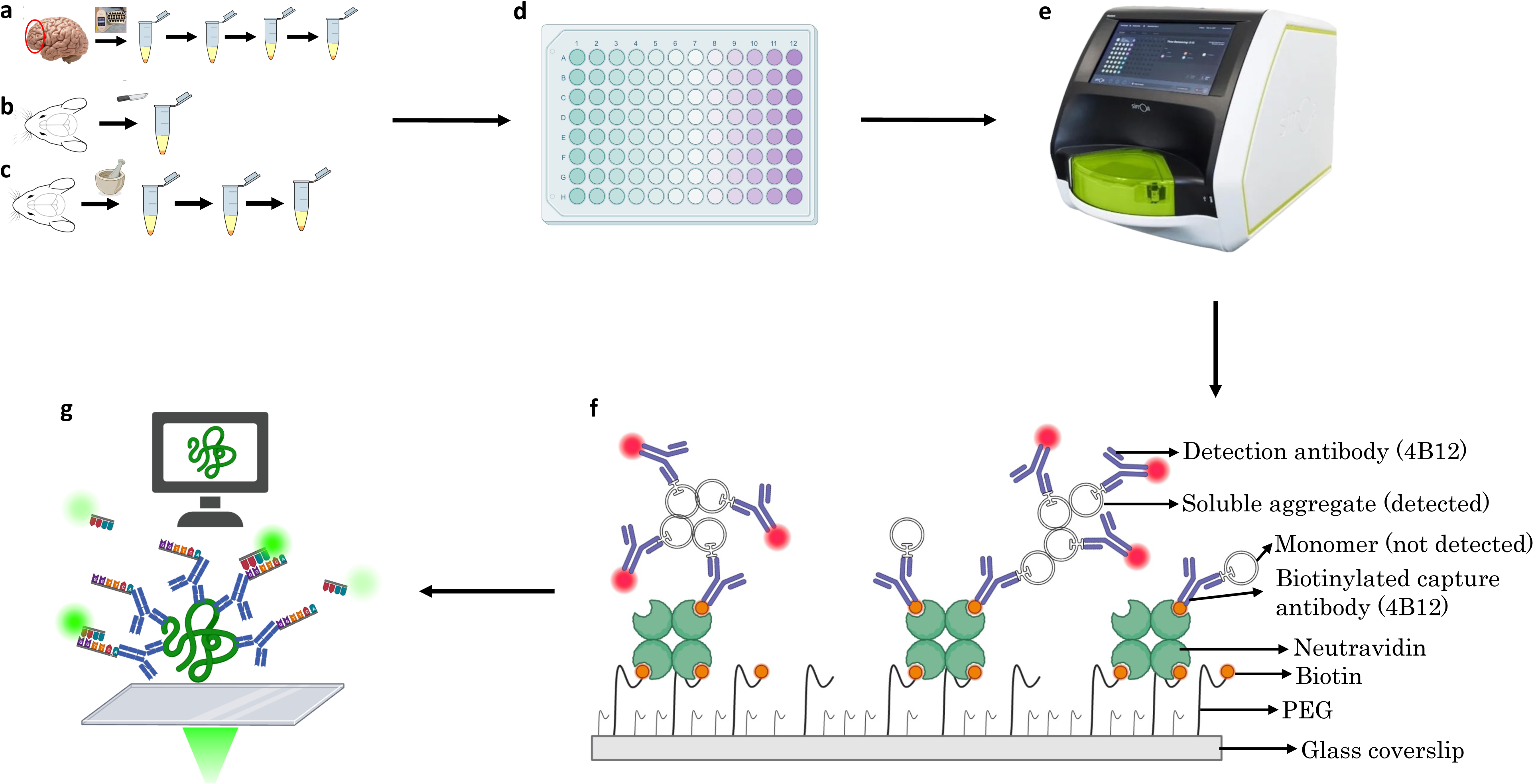
Graphic summary of the experimental design. Post-mortem human brain samples (**a**) along with Line 61 mouse brains (**b,c**) were processed by gentle soaking, detergent-free pulverised tissue homogenising and from which pellets then sequentially extracted using Triton X-100, and then sarkosyl. Once the soluble aggregates for the respective extraction methods were harvested, the total protein concentration in each sample was measured using a BCA assay (**d**), and the concentration of alpha-synuclein aggregates in the samples were determined using SIMOA (**e**). Then, the size and shape of the alpha-synuclein aggregates were characterised by SiMPull (**f**) and DNA-PAINT imaging (**g**).

**Figure 2.**
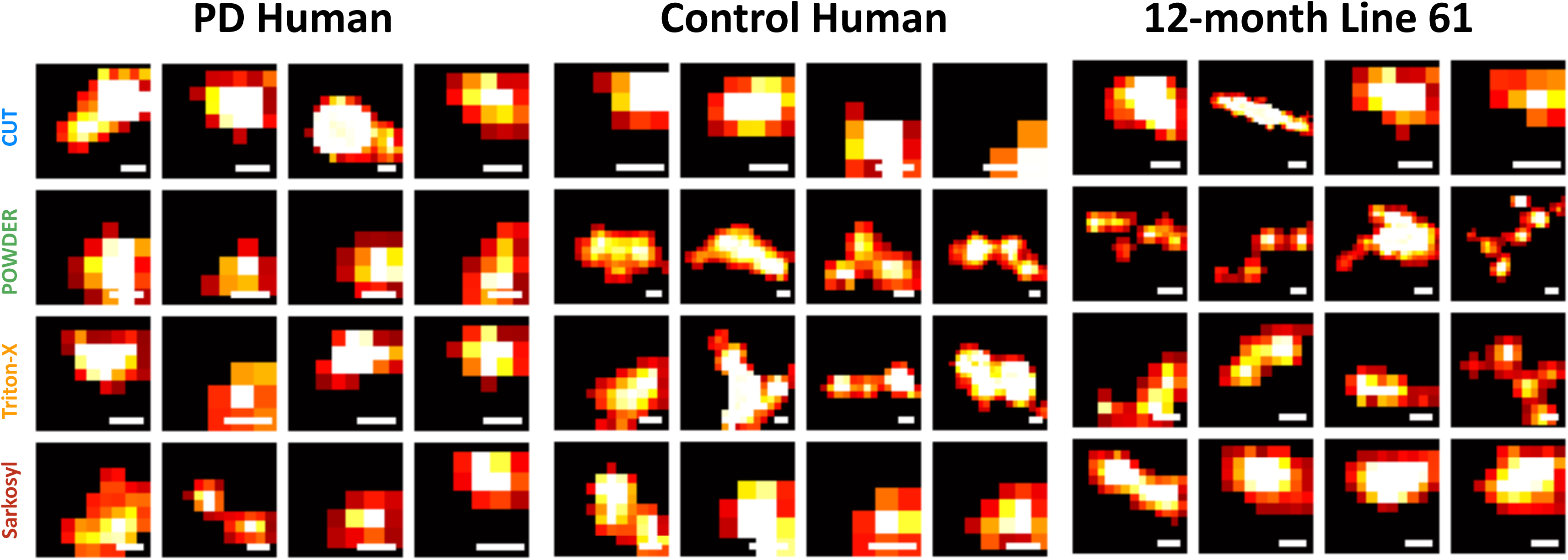
Representative micrographs of super-resolved aggregates PD (**a**) and control (**b**) post-mortem human, along with Line 61 mouse model at 12 months of age (**c**), processed by soaking (cut), homogenising (powder), Triton X-100, and sarkosyl extraction (scale bars are 20 nm). These randomly-selected aggregate images are provided as samples for the thousands of aggregates analysed ana averaged during the analysis.

**Figure 3.**
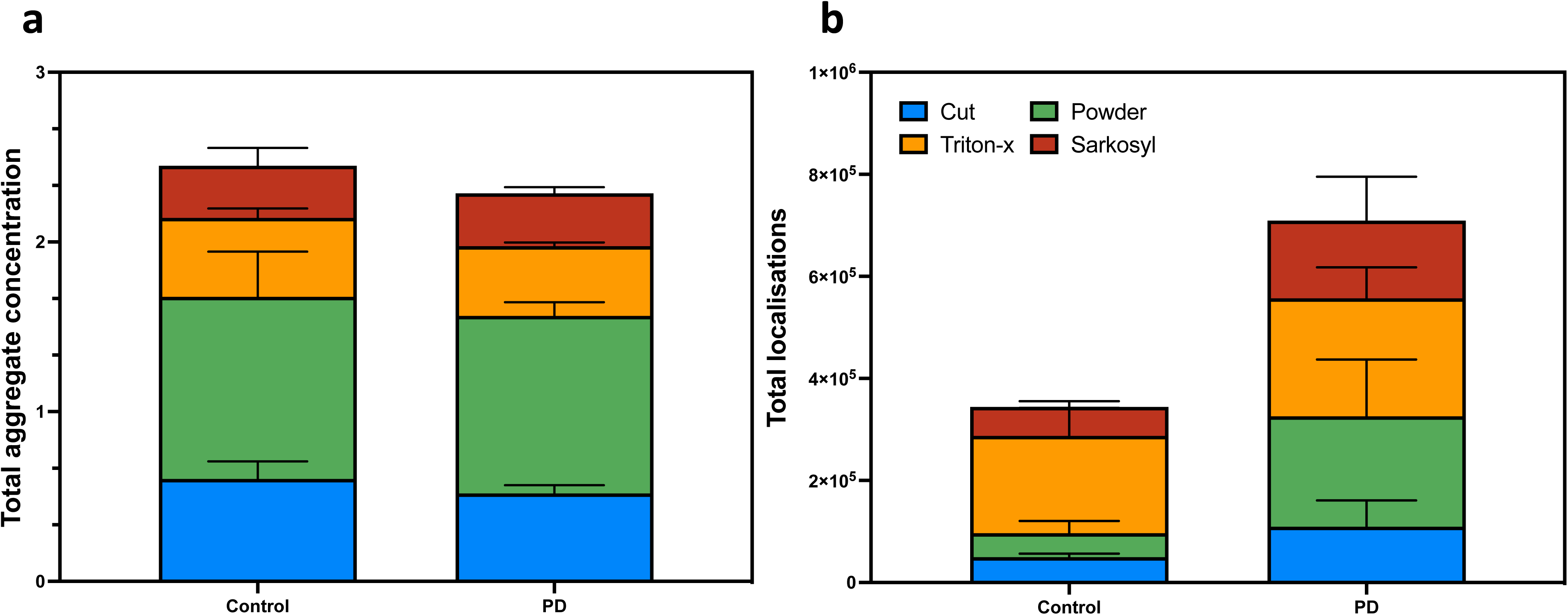
Total concentration of alpha-synuclein aggregates in the post-mortem human brain (**a**) samples measured by SIMOA for soaking (blue), homogenising (green), Triton X-100 (yellow), and sarkosyl (red) extractions. And total number of localisations measured by DNA-PAINT super-resolution microscopy (**b**).

**Figure 4.**
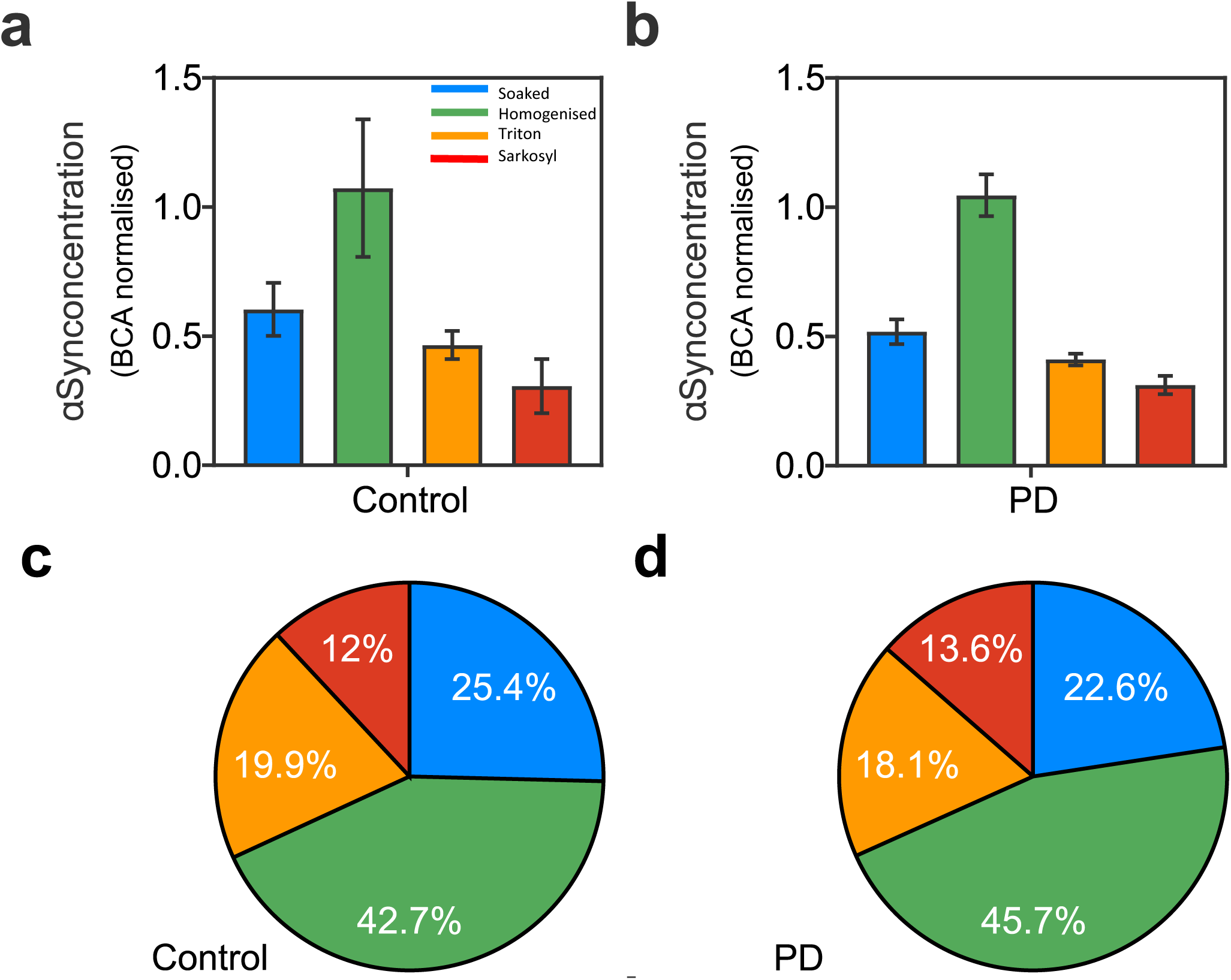
Concentration of the alpha-synuclein (□Syn) aggregates extracted by different methods from control (**a**) and PD (**b**) post-mortem human brain samples with SIMOA, and the proportion of the aggregates extracted by the serial processing from the control (**c**) and PD samples (**d**).

### 3.2 PD brains contain larger aggregates than controls

While the concentration of □Syn aggregates did not differ between the PD and control orbitofrontal cortex samples, morphology of these small aggregates differed significantly. PD brains contained longer (mean of 80 vs 65 nm) and larger (mean of 1360 vs 770 nm^2^) aggregates, with rounder (less fibrillar) shapes, regardless of extraction method, leading to higher total mass of aggregated □Syn in the PD brains (**Figure 3b**).

Aggregate morphology showed further differences based on the extraction method. From the control brains, the longest (mean of 74 and 70 nm; **Figure 5a**), largest (mean of 1140 and 835 nm^2^; **Figure 5b**), and roundest (mean of eccentricity 0.83 and 0.83; **Figure 5c**) aggregates were harvested by homogenising and by sarkosyl extraction. On the other hand, the homogenised and TrX extracted aggregates were longer (mean of 93 and 90 nm; **Figure 5d**) and larger (1980 and 1490 nm^2^; **Figure 5e**), from the PD brains, while the shortest (mean of 63 nm) and smallest (mean of 950 nm^2^) aggregates were in the sarkosyl extracted samples. Unlike the control brains, the larger aggregates were more fibrillar in the PD brain, as the aggregates with the highest eccentricity were found in the homogenised fraction, followed by the TrX extraction, and the roundest aggregates were in the sarkosyl soluble fraction (**Figure 5f**).

**Figure 5.**
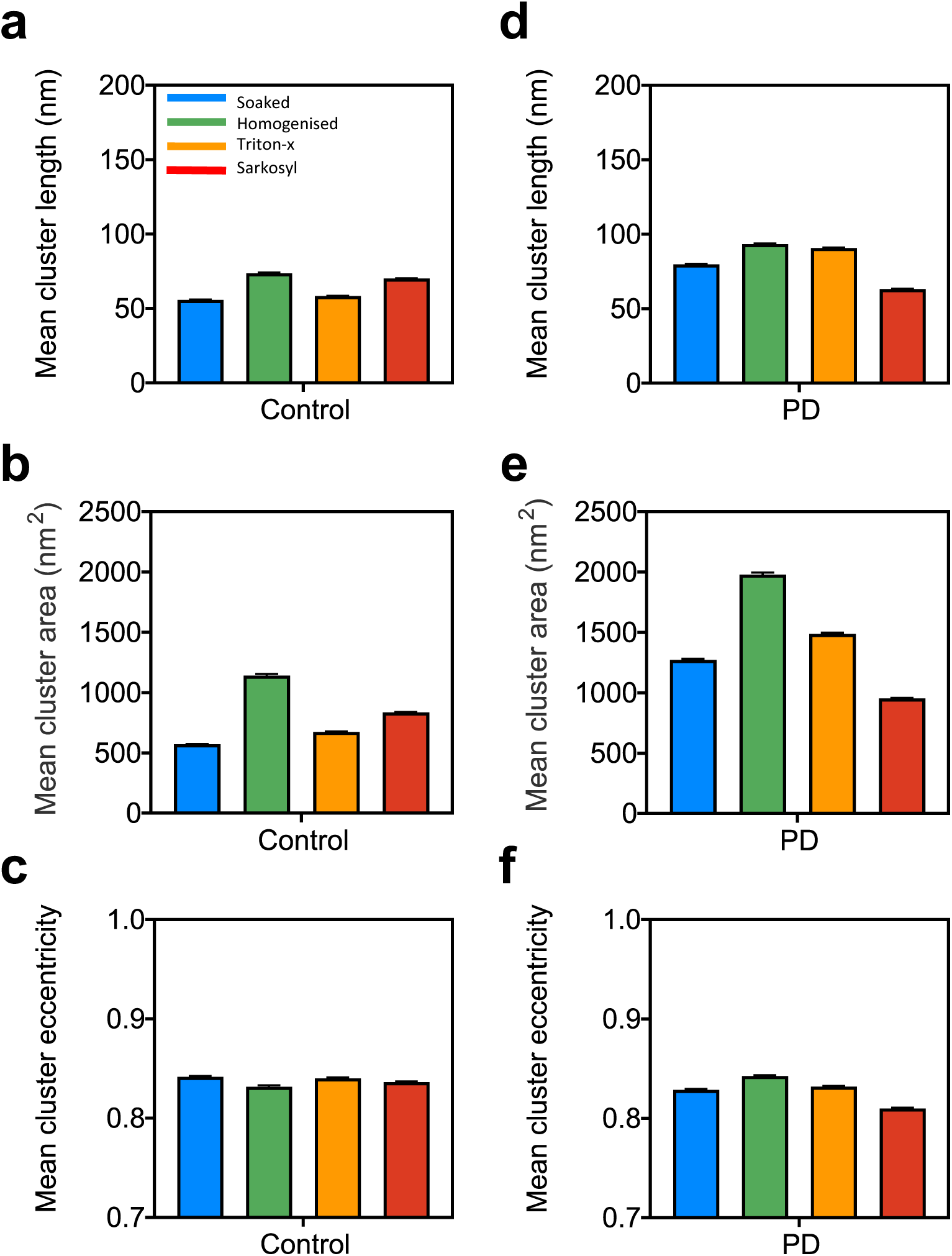
Mean alpha-synuclein aggregate cluster length (nm), area (nm^2^), and eccentricity for aggregates harvested from the control (**a, b, c**, respectively) and PD (**d, e, f**, respectively) post-mortem human brain samples harvested by gentle soaking (blue), homogenising (green), Triton X-100 (yellow) and sarkosyl (red) extractions.

We have recently developed a mathematical model which allows us to interpret aggregate size distributions in terms of the underlying molecular mechanisms(25). Briefly, for aggregates well above the nucleation size, our model predicts a geometric decay of the aggregate abundance with size. The rate of this decay is determined by the relative rates of aggregate formation and removal. When a cell is in a runaway aggregation, or “diseased” cell state, its rate of aggregate accumulation relative to removal is increased considerably, leading to a longer length distribution. In samples from PD cases, we expect some cells to still be “healthy”, producing shorter aggregates, and some cells to be “diseased” producing larger aggregates. We quantified the balance of production and removal of aggregates in cells as the relative removal rate. This analysis of the size distribution yields three quantities: the relative removal rate in heathy (1) and diseased (2) cells, along with the fraction of aggregates originating from diseased cells (3).

In this experiment, we found that the relative removal rates in healthy (1) and diseased cells (2) are conserved across samples from both control and PD brains. This implies that the aggregation behaviour of the healthy and diseased cell states are the same across PD and healthy aging. Meanwhile, the main difference between PD and control brains is that the amount of aggregates from the diseased cells is significantly higher in PD, most likely due to the increased number of diseased cells in PD. Moreover, we found differences in the results depending on the extraction method: while the relative removal rate between the healthy and diseased states is comparable between the soaked, homogenised, and TrX-extracted samples, with the relative removal in diseased cells being only about 30% that of the relative removal in healthy cells, for sarkosyl the difference is only approximately 50%, making sarkosyl an outlier (**Figure 6c**). As such, we excluded the sarkosyl-extracted samples from the analysis of fraction of aggregates that came from diseased cells. For the other three extraction methods, when we compared the fractions between the PD and control cases, the greatest difference was observed in the soaked samples, followed by TrX, meanwhile the homogenised samples showed the smallest difference (**Figure 6d**).

**Figure 6.**
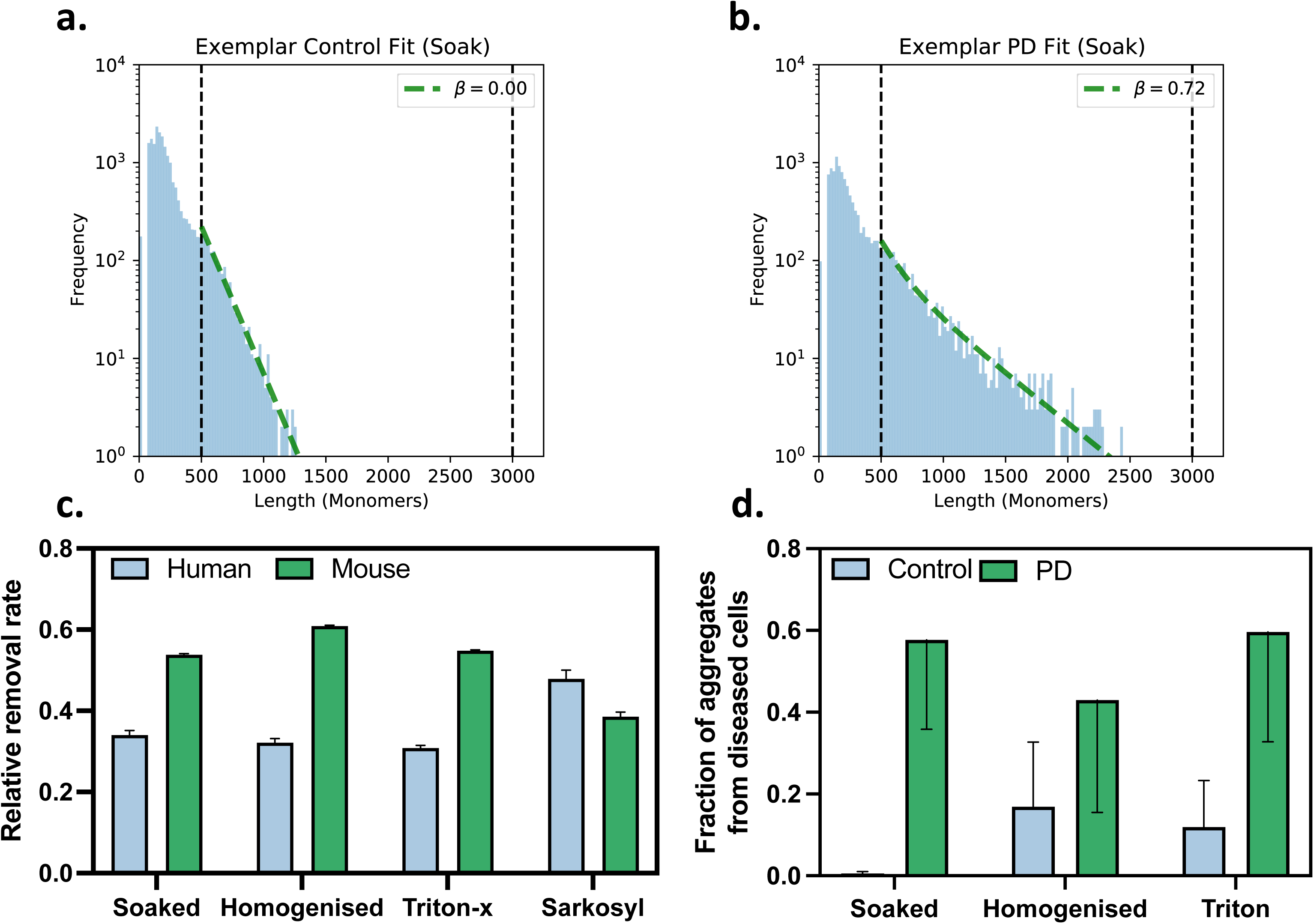
Example histograms of the aggregate size distribution for control (**a**) and PD (**b**) samples with the soak extraction. The green dashed line shows the predicted distribution using the model and the mean parameters determined from the Bayesian inference. The difference in the mean β from the fitting procedure reflects the difference in the size of the disease subpopulation between the control and PD samples. This can also be seen directly from the longer tail in the PD dataset. The relative removal rate in diseased cells compared to healthy cells (**c**). For human samples from soaked, homogenised and TrX extraction methods, the relative removal rate in diseased cells is decreased to 30% of the relative removal rate in healthy cells, consistent across the 3 extraction methods, indicating a considerable difference between the healthy and diseased states. For the mouse samples, and the sarkosyl extraction for the human samples, this difference is less pronounced, with the relative removal rate in diseased cells being only about half of that in healthy cells. In samples from control brains, the majority of aggregates comes from healthy cells, with the contribution from diseased cells being essentially undetectable in the soaked samples. By contrast, in samples from individuals with PD, the majority of aggregates came from diseased cells (**d**).

### 3.3 □Syn aggregate concentration in Line 61 mouse brain

Following the human samples, we quantified the small □Syn aggregate concentration in the 1.5-, 6-, 9-, and 12-month-old Line 61 mouse brain samples extracted by the same methods, in order to compare the aggregates produced in the human brains, with a mouse model expressing human □Syn at moderately high levels. While the □Syn aggregate quantity did not differ between the ages, concentration of aggregates harvested by homogenising, TrX, and sarkosyl extractions differed significantly (**Figures 7a-d**). Similar to the human PD cases, the highest concentration of □Syn aggregates was found in the homogenised samples, followed by the TrX soluble extract, and the sarkosyl soluble fraction (**Figures 7e-h**). Meanwhile, the soaked samples, which were prepared from a different brain sample contained a smaller amount of aggregates, compared to the other extraction methods.

**Figure 7.**
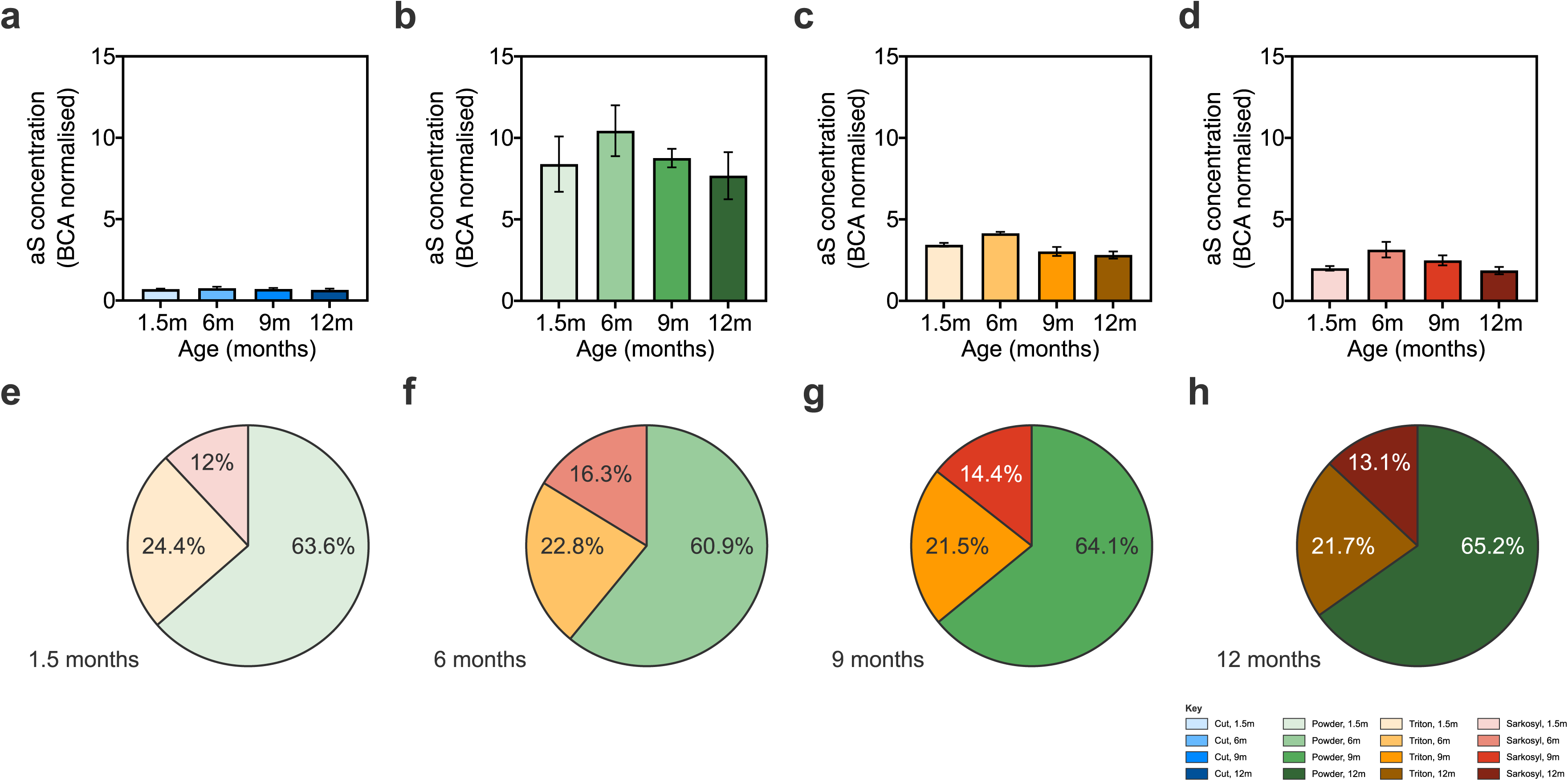
Concentration of alpha-synuclein aggregates in the Line 61 mouse brain measured by SIMOA for soaked (**a**), homogenised (**b**), Triton X-100 (**c**), and sarkosyl (**d**) extracted samples and the proportion of the aggregates extracted by the serial processing at 1.5- (**e**), 6-(**f**), 9- (**g**), and 12-months (**h**) of age.

For the soaked samples, the average aggregate length was 100 nm at 1.5-months of age and increased to 114 and 123 nm by 6- and 9-months of age but decreased to an average of 74 nm at 12-months of age, suggesting a change in the morphology of soaked aggregates as the Line 61 mice age (**Figures 8a & e**). Aggregate area from the soaked samples followed a similar trend, with an increase at 9-months of age, followed by a decrease at 12-months (**Figures 8a & i**). Meanwhile, aggregate eccentricity did not change with age and had a value close to 1, indicating the presence of more fibrillar aggregates in all samples (**Figures 8a & m**). Within the serially extracted samples, the homogenised and TrX soluble aggregates had an average length of 128 and 142 nm respectively, and thus were longer than the sarkosyl soluble aggregates, which had an average length of 59 nm. However, while the TrX soluble aggregates grew longer as the mice aged, the homogenised aggregates varied in length and showed no clear trend with age. In contrast, while the sarkosyl soluble aggregates grew longer after 1.5-months of age, they did not show any further significant age-related changes (**Figures 8b-d & f-h**). The area of the TrX soluble aggregates grew larger with age, while homogenised and sarkosyl soluble samples had a fluctuating pattern (**Figures 8b-d & j-l**). On the other hand, eccentricity of the serially extracted samples differed by extraction method. While the homogenised and TrX soluble aggregates were more fibrillar, the sarkosyl soluble aggregates were more circular (**Figures 8b-d & n-p**).

**Figure 8.**
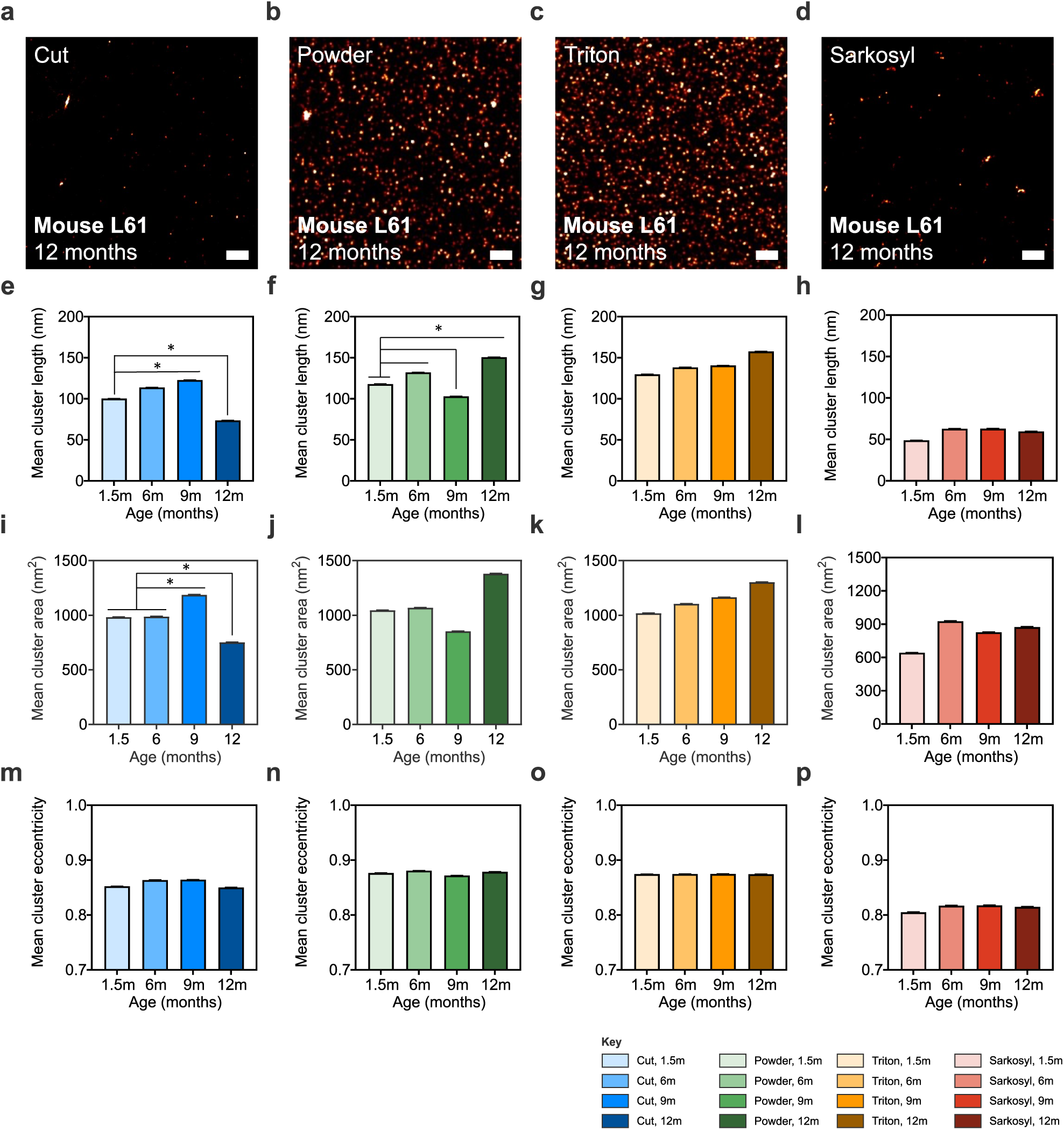
Representative micrographs of super-resolved aggregates from the 12-month-old, Line 61 mouse brain processed by soaking (**a**), homogenising (**b**), Triton X-100 (**c**), and sarkosyl (**d**) extraction (scale bars are 500 nm). Mean alpha-synuclein aggregate cluster length (nm), area (nm^2^), and eccentricity for aggregates harvested from the Line 61 mouse brain by soaking (**e, i, m**, respectively), homogenising (**f, j, n**, respectively), Triton X-100 (**g, k, o**, respectively), and sarkosyl (**h, l**, **p**, respectively) extraction.

### 3.4 Comparing PD human and transgenic mouse aggregates

An essential question in the field of neurodegeneration and in the broader field of neuroscience is the validity of animal models. Since the morphology of pathological aggregates is related to their toxic properties(16,30), an important requirement of a useful mouse model for studying disease pathogenesis and testing novel therapeutics is likely to be that the aggregates formed are similar to the ones in humans. In order to evaluate the Line 61 mouse model of PD in terms of its aggregate validity, we once again employed our mathematical model. In all extraction methods, except sarkosyl, the mouse data showed significantly less difference between the healthy and the diseased cell states than the human data, suggesting a decrease in the relative rate of removal of only 50 – 60% (**Figure 6c**). In other words, while there is clear evidence for a healthy and a diseased cell population in both PD and control human brains, in the mouse, the distinction between two such populations is less clear. Furthermore, the length distributions observed in the mouse samples suggest that the relative removal rate in mouse cells corresponds to neither the healthy nor the diseased cell state in humans. For soaked and homogenised samples, the mouse rates lie between the healthy and diseased values for human samples. Meanwhile, for TrX, the relative removal rate in the mouse is always lower.

## Discussion

In this study, we quantified and characterised nanoscopic □Syn aggregates from human post-mortem PD and control orbitofrontal cortex samples, along with the commonly used Line 61 mouse model of PD. We processed the tissue using serial extractions with four commonly used methods to harvest different types of nanoscopic aggregates. Gentle soaking was used initially to harvest diffusible aggregates, followed by detergent-free homogenisation to solubilise the aggregates that are not diffusible but also not membrane bound. Then the tissue samples were further treated with TrX to digest the cellular membrane and solubilise the membrane bound aggregates, followed by a sarkosyl extraction, which partially solubilises the fibrillar aggregates. As such, we were able to address five questions: (1) how do the □Syn aggregates from post-mortem PD brain samples compare to controls?; (2) do different extraction methods harvest morphologically different types of aggregates?; (3) does the quantity or morphology of the □Syn aggregates change as the Line 61 mice age?; (4) how do the □Syn aggregates from a mouse model compare to human samples?; and (5) can the aggregate size distribution be fitted by chemical kinetics and what does this reveal about the aggregation mechanisms of □Syn *in-vivo*?

The human post-mortem tissue samples were selected from the orbitofrontal cortex at Braak stages 4 and 5. Since this region has extensive dopaminergic input from substantia nigra, executive dysfunction is one of the major symptoms of PD(31) and the orbitofrontal cortex volume and pathology is linked to deficits in reward processing and decision making(32,33). In this experiment, we used orbitofrontal cortex samples at Braak stages 4 and 5, which is prior to Lewy body formation in this region, in order to characterise the nanoscopic aggregates in a disease-relevant region, before significant Lewy body pathology and neuronal loss (34).

Remarkably, the aggregate numbers did not differ between PD and control brains as the orbitofrontal cortex samples from non-PD brains also contained small aggregates at comparable concentrations to PD brains. However, the aggregates in the PD brain samples were significantly larger, leading to a greater total aggregated □Syn mass. This suggests that even in a region without Lewy body pathology at the studied stage, morphological differences in nanoscopic aggregates exist in disease, indicating that there has already been a disruption of protein homeostasis and hence increased aggregation. However, if all the neurons showed increased aggregation, then one would expect an increase in aggregate concentration along with an increase in average aggregate size. On the other hand, if only a fraction of neurons have disrupted protein homeostasis leading to growth of longer aggregates, this could result in an increase in average length but no change in aggregate concentration. Indeed, fitting the data to our mathematical model showed that the fraction of aggregates from cells with disrupted aggregate removal mechanisms increased from low to undetectable levels in control samples to the majority in samples from PD. This effect was also dependent on the extraction method, with the most pronounced differences in aggregates from soaking and TrX extraction. On the other hand, even though the greatest quantity of aggregates were in the homogenised samples, the fraction of aggregates that came from diseased cells showed the smallest difference between the PD and control cases for this method.

Both the concentration and morphology of the aggregates differed significantly between the tissue harvesting methods. The least amount of aggregates from both the PD and control brains were harvested by sarkosyl extraction, which were also the shortest aggregates in the disease samples. Since sarkosyl partially solubilises fibrillar aggregates, the lack of Lewy body pathology and in parallel small amount of insoluble (fibrillar) aggregates in the orbitofrontal cortex at this disease stage explains the low quantity of sarkosyl-soluble aggregates in the samples. Importantly, our kinetic model also showed the least mechanistic information was contained in sarkosyl extracted aggregates, as judged by the similarity between diseased healthy and mouse samples. This suggests that the size distributions obtained by sarkosyl extraction are dominated by the effects of the harsh extraction conditions that disrupt the morphology of the aggregates, rather than reflecting the size distribution that existed *in-situ*. Meanwhile, the soaked and TrX-soluble aggregates showed a higher disease-related signal than the homogenised samples even though their concentrations were not as high. As shown previously, the diffusible aggregates harvested by gentle soaking may have higher toxic properties(16,17), contributing to pathogenesis of PD. Similarly, it has been shown that □Syn aggregates may grow on the lipid membrane(20,22) and as such, these TrX-harvested aggregates, which would expected to be membrane associated, may be more relevant to early PD pathology.

Our previous work imaged the aggregates in the soaked fraction in the amygdala, which unlike the orbitofrontal cortex shows high levels of pathology and neuronal loss at this stage(35). This study used a different super-resolution method based on an aptamer that imaged beta-sheet containing aggregates (both beta-amyloid and □Syn), thus the results cannot be directly compared with the current study. Nevertheless, it is noteworthy that we have shown smaller aggregates in the PD samples compared to the controls, suggesting that prior to neuronal loss, larger □Syn aggregates form (as seen in the orbitofrontal cortex), which then possibly leads to the loss of the neurons containing these larger aggregates, leaving the cells with smaller aggregates behind. This also highlights that in future work it will be informative to compare the aggregates in different brain regions at different disease stages using the same imaging method, as PD pathology seems to progress at different rates in different brain regions, with the formation of larger aggregates leading to cell death and related behavioral phenotypes.

Following the characterization of human brain-derived aggregates, we also compared them to them Line 61 mouse model of PD at different ages. Histopathological □Syn accumulation starts in the Line 61 mice by 1-month of age(36). As demonstrated by Rabl *et al*.,(37) while the total (monomeric and aggregate) TrX soluble human □Syn levels increase until 6-months of age in the hippocampus, they decrease in the striatum. On the other hand, the TrX insoluble fragments does not change significantly in either region(37). By different methods, our results show that the number of aggregates does not change over time, yet their size increases with age, leading to greater total □Syn mass in the brains of older mice. As such, the constant presence and slow growth of the □Syn aggregates may eventually lead to PD-like pathology. However, this age-dependent increase in aggregate size was not detectable with all extraction methods; while the aggregates in the homogenised and TrX soluble fragments grew larger by 12-months of age, the aggregates in the soaked samples got smaller. This harvesting method-dependent difference in aggregate morphology demonstrates the presence of aggregates accessible by different methods in the brain tissue and highlights the importance of studying the relationship between the harvesting method and the aggregates collected.

Compared to the human brain samples, the Line 61 mice had a significantly higher (~10-fold) aggregate load compared to the humans, probably due to the over-expression of the *SNCA* gene. We have also found that the □Syn aggregates in the human brain are shorter with larger total areas, and rounder compared to the ones in the Line 61 mouse brain. While the average length of human aggregates was under 100 nm, agreeing with previous findings from our group(16), the aggregates in the mouse brain - especially the ones in the homogenised and TrX soluble fragments were about 150 nm in length. The presence of longer aggregates in the mice may be due to the over-expression of □Syn, meaning that the aggregate removal processes are overwhelmed in a larger number of cells, resulting in more aggregates than in human. Humans have less rapid aggregation and hence removal processes are more effective, so that fewer larger fibrillar aggregates form. Overall, the proportion of aggregates in the different fractions were remarkably similar between the humans and the Line 61, suggesting that the aggregate properties which determine the relative amounts in each fraction are similar in both animals and humans. Nevertheless, the morphological differences between the human disease samples and the mouse model are concerning. Since the small aggregates can interact with different cellular receptors and other proteins, giving rise to various disease mechanisms, the differences between the morphology of these aggregates reduces the validity of this model. Indeed, the lack of dopaminergic neuronal loss and heavier pathology seen in the hippocampus, may be at least partly due to these differences in small aggregate characteristics, which should be considered while working with this mouse model.

## Conclusion

Our results show that different populations of □Syn aggregates exist in different sub-populations of neurons within a brain region, which can be harvested using different methods. Importantly, the diseased and healthy cells in the PD and non-PD brains behave similarly biologically, yet their proportion is increased in PD. While nanoscopic aggregates are also produced in non-PD brains at similar rates to PD, the clearance of these aggregates is reduced in PD, leading to further growth of these aggregates, increasing the total aggregated □Syn mass. This may eventually lead to the formation of Lewy bodies in the PD brains, promoting cell death. A such, simply quantifying these aggregates is not sufficient to gain insights on PD mechanisms and advanced single-molecule imaging methods are required to characterize these nanoscopic aggregates individually, which can then be analysed using mathematical models, to understand the pathogenesis of PD.

## List of abbreviations

□Syn: Alpha synuclein
aCSF: Artificial cerebrospinal fluid
DLB: Dementia with Lewy body
DNA-PAINT: DNA Point Accumulation in Nanoscale Topography
FRET: Fluorescence energy transfer
PD: Parkinson’s disease
Sarkosyl: N-Laurylsacosine
SIMOA: Single-molecule array
SiMPull: Single-molecule pulldown
SNARE: Soluble N-ethylmaleimide-sensitive factor attachment protein receptor
TBS: Tris-buffered saline
Thy1: Thymocyte differentiation antigen 1
TIRF: Total internal reflection fluorescence
TrX: Triton X-100

## Resource availability

### Lead contact

Requests for further information and resources should be directed to Professor Sir David Klenerman, Yusuf Hamied Department of Chemistry, University of Cambridge.

### Materials availability

This study did not generate new unique reagents.

### Data and code availability

- Data availability: Data reported in this paper will be shared by the lead contact upon request.
- Code availability: All original code has been deposited at github and is publicly available as of the date of publication. DOIs are listed in the key resources table
- Any additional information required to analyse the data reported in this paper is available from the lead contact upon request.

## Declarations

### Ethics approval and consent to participate

Human post-mortem brain tissue was acquired from the Cambridge Brain Bank (Cambridge University Hospitals). The Cambridge Brain Bank is supported by the NIHR Cambridge Biomedical Research Centre (NIHR203312). We gratefully acknowledge the participation of all our patient and control volunteers.

### Consent for publication

Not applicable.

### Availability of data and materials

Data collected during these experiments will be made available upon reasonable request from the corresponding author. No novel code was generated during data analysis.

### Competing interests

None to declare.

### Funding

The work was supported by the UK Dementia Research Institute (which receives its funding from UK DRI Ltd), the UK Medical Research Council, Alzheimer’s Society and Alzheimer’s Research UK (ARUK-PG2020A-009), ARUK-PG2020A-009, the Royal Society and a grant from Eisai. The views expressed are those of the authors and not necessarily those of the NHS, the NIHR or the Department of Health.

### Author contributions

E.F. Conception and design, data collection and statistical analysis, manuscript writing. J.S.H.D. Conception and design, data collection and image analysis, manuscript editing. S.N. Conception and design, data collection, manuscript editing. G.M. Model building and data fitting. M.B., J.Y.L.L., Z.X., Y.W., B.P., Data collection, Y.I. Conception and design. A.Q. Human brain sample preparation and characterisation. J.S. Conception and design, manuscript editing. D.K. Conception and design, manuscript editing, overall supervision of the project.

## Acknowledgments

We would like to thank Dr. Rohan T. Ranasinghe for his help during data collection.

## Figure captions

**Supplemental Figure 1.** Histograms of the measured aggregate length distributions in human samples for different extraction techniques. The red dashed line shows the predicted distribution using the model and the mean parameters determined from the Bayesian inference.

**Supplemental Figure 2.** Posterior distributions for the fraction of aggregates from diseased cells following the Bayesian inference fitting procedure. The dashed black line shows the mean value for each experiment.

**Supplemental Figure 3.** Histograms of the measured aggregate length distributions in mouse samples for soaked (cut) samples. The red dashed line shows the predicted distribution using the model and the mean parameters determined from the Bayesian inference.

**Supplemental Figure 4.** Histograms of the measured aggregate length distributions in mouse samples for homogenised (powder) samples. The red dashed line shows the predicted distribution using the model and the mean parameters determined from the Bayesian inference.

**Supplemental Figure 5.** Histograms of the measured aggregate length distributions in mouse samples for Triton X-100 extracted samples. The red dashed line shows the predicted distribution using the model and the mean parameters determined from the Bayesian inference.

**Supplemental Figure 6.** Histograms of the measured aggregate length distributions in mouse samples for sarkosyl extracted samples. The red dashed line shows the predicted distribution using the model and the mean parameters determined from the Bayesian inference.

## Methods

### EXPERIMENTAL MODEL AND STUDY PARTICIPANT DETAILS

3 PD, and 3 age- and sex-matched (two female, one male) post-mortem human orbitofrontal cortex (Brodmann areas 10-11) samples were used alongside brain samples from Line 61 mice. The human disease brain samples were selected from Lewy body Braak stages 4/5 (38). Due to the extensive dopaminergic input to this region from the midbrain, the orbitofrontal cortex is involved in the decision making and reward processing deficits seen in PD(32,33). Brain samples were snap frozen and stored frozen at –80□ until synaptosome preparation. Human post-mortem brain tissue was acquired from the Cambridge Brain Bank (Cambridge University Hospitals). The Cambridge Brain Bank is supported by the NIHR Cambridge Biomedical Research Centre (NIHR203312). We gratefully acknowledge the participation of all our patient and control volunteers.

The Line 61 model was first developed and described by Rockenstein *et al.*(39), as a X chromosome-linked transgenic model, expressing human *SNCA* gene under the murine thymocyte differentiation antigen 1 (Thy1) promoter in C57BL/6xDBA/2 mice(40). Unlike many mouse models over-expressing disease-associated proteins at unphysiologically high levels, the Line 61 mice express full-length human □Syn at biologically-relevant levels throughout the brain; histopathological and Western blot analysis showed a 1.5- to 3.4-fold rise in expression(36), which is comparable to individuals with familial PD due to *SNCA* gene triplication(41). While □Syn accumulation starts around 1 month of age, no significant dopaminergic or motor neuron loss has been reported, even at advanced ages(36,39). On the other hand, hippocampal neuronal loss - especially in the CA3, is apparent by 3 months of age(42) and deficits in dopamine release and striatal pathology, along with increased gliosis (both astrocytes and microglia), and mitochondrial dysfunction are observed by 6-months of age(43–46). Motor and non-motor behavioural dysfunction has also been reported in this model, starting as early as 1-month and progressing with age(36,37).

Due to the appearance of □Syn accumulation by 1-month and the pathophysiological symptoms by 6-months of age, and the age-related symptom progression, 1.5-, 6-, 9-, and 12-month-old mouse brains were used in this study, acquired from QPS (now Scantox Neuro). Brains were harvested from a total of 24 male Line 61 (6 mice per group at 1.5-, 6-, 9- and 12-months of age), following euthanasia with sodium phenobarbital (200 mg/kg) and perfusion with phosphate buffered saline (PBS 10%). 12 of the brains were used for the gentle soaking extraction (**Figure 1b**), while the remaining 12 were powdered (homogenised) and serial extractions with artificial cerebrospinal fluid (aCSF), TrX, and sarkosyl were performed (**Figure 1c**). While the lack of female mice is a limitation of this work, this decision was based on the X chromosome-linked expression of *SNCA* in this model, as the stochastic inactivation of the X chromosome may lead to differences of expression between neurons and confound the results.

### METHOD DETAILS

#### Human tissue collection and sample preparation

Post-mortem orbitofrontal cortex samples were first dissected into 300-350 mg pieces (**Table 1**). For the gentle soaking protocol, tissue samples were first cut into small pieces with a scalpel and transferred to 1.5 mL protein low-binding Eppendorf tubes (Eppendorf, Cat. 0030108116). 600 µL of aCSF buffer (bio-techne TORIS, Cat. 3525) with Halt^™^ protease and phosphatase inhibitor single-use cocktail (100x; Thermo Fischer Scientific, Cat. 78442) was added to each sample and incubated for 45-minutes at 4□ on a HulaMixer^™□^ (Thermo Fischer Scientific, Cat. 15920D). Then the samples were centrifuged at 17,000 G for 120-minutes at 4°C, the supernatant was collected and stored in a −80°C freezer until analysis. Subsequently, the pellet was resuspended with 500 µL aCSF with protease/phosphatase inhibitors and 1 mm zirconium beads (Scientific Labs, Cat. SLS1414) were added. Then the samples were processed on an electronic tissue homogeniser (VelociRuptor V2 Microtube Homogeniser, Scientific Labs, Cat. SLS1401), at 5 meters/sec for 2 cycles of 15 seconds, with a 10 second gap in between, followed by centrifugation at 17,000 G for 120-minutes at 4□. The supernatant was collected and stored in a −80°C freezer until analysis as the “homogenised” sample. Meanwhile, the centrifugation pellet was resuspended with 500 µL aCSF containing inhibitors as described above, as well as 1% (by volume) TrX (Fischer Scientific, Cat. 11488696) and incubated for 45-minutes at 4□ on a HulaMixer^™□^. Then the samples were centrifuged at 17,000 G for 120-minutes at 4°C, the supernatant (TrX soluble sample) was collected and stored in a −80°C freezer until analysis. Then the centrifugation pellet was once again resuspended with 500 µL aCSF containing inhibitors as described above, as well as 1% (by volume) sarkosyl (Sigma-Aldrich, Cat. 61747-100ML) and the same extraction described above was performed to acquire the sarkosyl soluble sample (**Figure 1a**).

#### Mouse sample preparation

For the gentle soaking protocol, mouse cerebral tissue was first cut down into small pieces with a scalpel and transferred to 1.5 mL lo-binding Eppendorf tubes. 600 µL of aCSF buffer with Halt^™^ protease and phosphatase inhibitor single-use cocktail was added to each sample and incubated for 45-minutes at 4□ on a HulaMixer^™□^. Then the samples were centrifuged at 17,000 G for 120-minutes at 4°C, the supernatant was collected and stored at −80°C until analysis.

For the serial extractions, the intact mouse cerebrum was first frozen with liquid nitrogen and grounded into powder using a mortar and pestle. Then the powder was transferred into a 1.5 mL lo-binding Eppendorf tube and the initial aCSF extraction was performed as described above. The supernatant (homogenised/powder sample) was collected and stored at −80°C until analysis. Subsequently, the pellets were further extracted with TrX and sarkosyl, in the same way as described for the human samples.

For the serial extractions, the intact mouse cerebrum was first frozen with liquid nitrogen and grounded into powder using a mortar and pestle. Then the powder was transferred into a 1.5 mL protein low-binding Eppendorf tube and the initial aCSF extraction was performed as described above. The supernatant (homogenised/powder sample) was collected and stored at −80°C until analysis. Meanwhile, the centrifugation pellet was resuspended with 500 µL aCSF containing inhibitors as described above, as well as 1% (by volume) TrX and incubated for 45-minutes at 4□ on a HulaMixer^™□^. Then the samples were centrifuged at 17,000 G for 120-minutes at 4°C, the supernatant (TrX soluble sample) was collected and stored in a −80°C freezer until analysis. Then the centrifugation pellet was once again resuspended with 500 µL aCSF containing inhibitors as described above, as well as 1% (by volume) sarkosyl and the same extraction described above was performed to acquire the sarkosyl soluble sample.

#### BCA assay

Total protein concentration in the samples processed by different methods was determined using a bicinchoninic acid (BCA) assay (**Figure 1d**; Pierce^™^ BCA Protein Assay Kit, Thermo Fischer Scientific, Cat. 23227). In brief, samples were diluted in PBS (1:5 by volume) and 25 µl diluted sample was loaded in duplicates into a 96-well plate. Bovine serum albumin (BSA) assay standards were also diluted in PBS at instructed amounts by the kit and loaded alongside the samples, also at 25 µl. Assay working reagent was prepared by mixing 50 parts of BCA Reagent A with 1 part of BCA Reagent B (50:1, Reagent A:B), provided with the kit and added at 200 µl to each sample. The plate was gently shaken for 30 seconds, covered, and incubated at room temperature for 30 minutes. Then the absorbance was measured at 562 nm on a plate reader.

#### SIMOA assay

□Syn aggregates in the mouse and human brain samples were quantified using an □Syn aggregate-specific single-molecule array (SIMOA; **Figure 1e**) we have developed(26). SiMoA^®^ Homebrew carboxylated beads (Quanterix; Cat.104006) were first functionalised with monoclonal 4B12 antibody (Thermo Fisher Scientific; Cat. MA1-90346) as described in the Quanterix Homebrew instruction manual. In brief, antibodies were buffer-exchanged with the provided bead conjugation buffer and 1 mL of the paramagnetic carboxylated beads (4.2×108 beads) were washed with the provided bead wash buffer followed by the bead conjugation buffer with the. 9 µL of EDC solution was added to the washed beads and the reaction mixture was incubated on a mixer for 30 minutes at 4□. After washing the activated beads once with cold 25 mM MES, the buffer-changed antibodies (0.2 mg/mL, 300 µL) were added. The reaction mixture was incubated on a mixer at 4°C for two hours. The conjugated beads were then washed with the provided bead wash buffer and blocked with the blocking buffer at room temperature for 40 minutes.

The beads coated with the 4B12 antibody were first washed three times with the provided bead diluent in the kit. Meanwhile, to each well on a conical bottom microplate (Quanterix), conditioned media samples were diluted with the lysate sample diluent (2:98) to give 100 µL solution at the desired concentration. Then, 25 µL of the washed beads were introduced to each well to give the final-bead concentration as 2 × 10^7^ beads/mL. The samples and the beads were incubated on the plate shaker at 30°C at 800 revolutions per minute (rpm) for 30 minutes. The plate was then washed by the SIMOA washer with the provided buffers. Then, the biotinylated detector antibody 4B12 was diluted to 0.1 µg/mL with the provided Homebrew detector diluent and 100 µL of the diluted detectors were introduced to each well. The mixture was then incubated on the plate shaker at 30°C with 800 rpm for 10 minutes, followed by another wash. Meanwhile, the provided streptavidin-β-galactosidase (SBG) concentrate was diluted to 50 pM with SBG diluent. 100 µL of the diluted SBG was added to each well and the mixture was incubated on the plate shaker at 30°C at 800 rpm for 10 minutes. Finally, the plate was washed again and loaded onto the SR-X^™^ Biomarker Detection System (Quanterix) together with the SIMOA disc, tips, and an RGP bottle.

#### DNA-PAINT with SiMPull imaging

□Syn aggregates in the mouse and human brain samples were characterised in terms of their size and shape using DNA point accumulation in nanoscale topography (DNA-PAINT) microscopy combined with single-molecule pulldown (SiMPull)(27). SiMPull coverslips were prepared as described previously(16,27) and kept in vacuumed containers at −20□ until used. At the beginning of each experiment, one vacuum box containing the coverslips was taken out of the freezer and placed inside a fume-hood to allow equalisation of the temperature. The coverslip was first coated with neutravidin (0.2 mg/mL) diluted in TBST(v/v) (0.05% Tween 20 Cat. P1379-25ML diluted in tris-buffered saline (TBS)) for 10-minutes. After removing the neutravidin solution, each well was washed with TBST (two cycles) and then with TBS with 1% Tween (one cycle)-hereon described as washing. Biotinylated 4B12 antibody was diluted to 10 nM in TBS containing 0.1 mg/ml bovine serum albumin (BSA; Thermo Scientific; Cat. 10829410) and incubated for 15 minutes. Then the wells were washed, and samples were incubated overnight at 4□. While the cut, powder, and sarkosyl extracted samples were tested neat, the TrX extracted sample was diluted in TBS 50:50 by volume. In order to minimise non-specific binding, a blocking step was performed after samples incubation, with solution containing 3 mg/ml BSA in TBS, incubated for 30 minutes. Then the DNA-labelled 4B12 antibody diluted to 2 nM in TBS containing 0.1 mg/ml bovine serum albumin was incubated for 30 minutes (labelling protocol explained by Fertan et al., 2023(27)). After the washing steps, TetraSpeck microspheres (1:7,000 in TBS, 10 µL, Thermo Scientific; Cat. T7279) were introduced to each well for 10 minutes. The TetraSpeck solution was then removed, followed by a single wash with TBST, and a second PDMS gasket (Merck, GBL-103250-10EA) was stacked on the coverslip before introducing 4 µL of imaging strand (TGGTGGT-cy3B; atdbio) in TBS. Finally, the coverslip was sealed with another coverslip on top of the second PDMS gasket (**Figure 1f**). The coverslip was imaged using a purpose built total internal reflection fluorescence (TIRF) microscope(47) using a 568-nm (green) excitation laser for 4,000 frames per field of view (FoV) at an exposure time of 100 ms (**Figure 1g**). As measured previously(47) using DNA origami structures(48), the spatial resolution of this setup is ~30 nanometres. Aggregates below this length were not included in the analyses.

### QUANTIFICATION AND STATISTICAL ANALYSIS

#### Data analyses

All BCA and SIMOA assays were run in duplicates, and the data were averaged. DNA-PAINT imaging was performed for 4 FoVs per well and two wells for each sample, from which the data were pooled. Super-resolution images were reconstructed, drift corrected, and analysed using a software developed by our group(49,50). The R Project Statistical Computing version 4.2.2 (2022-10-31)—“Innocent and Trusting” was used for all statistical analyses, the graphs were generated in GraphPad Prism 7.0a for Mac OS X and Inkscape, and the cartoon figures were created with BioRender.com. Linear mixed effects models, ANOVAs with Type 2 sums of squares, and Welch Two Sample t-tests were used to determine differences between the groups and corresponding 95% confidence intervals (CI) were reported. The datasets generated and analysed during the current study are available from the corresponding author on reasonable request.

#### Distribution fitting

For the detailed derivation of our models see Cotton et al(25). Briefly, we assume that at aggregated sizes well above the nucleus size, the number of aggregates at each size is determined by the rate at which they grow compared to the rate at which they are removed, through any process. This predicts a geometrically decaying size distribution, i.e. a straight line when the histograms of aggregate population versus size are plotted on a logarithmic plot (**Figures 6a-b**). We expect this to describe the situation within an individual cell. To analyse the aggregate size distributions from a brain sample, we need to consider the fact that it will contain aggregates from many different cells, some of which may be significantly more affected by disease / contain significantly more aggregates than others. At the very least, we expect there to be two cell states a healthy one and a disease associated one, in which the relative rates of aggregate production and removal, and thus the length distributions, differ. There is evidence that this binary state is indeed a good description(25,51). We therefore fit the data to a model that allows 2 different cell states: a healthy one in which aggregate production is slower compared to removal, determined by the parameter α*_H_*, and a diseased cell state in which aggregate production is faster compared to removal, determined by the parameter α*_D_*. The free parameters are the relative rates of aggregate removal in each cell state and the fraction of aggregates from the diseased state. To determine these parameters from data, we convert the length in nm to a length in terms of the number of proteins by multiplying by 4 (assuming a periodicity of 2 monomers per nm and a double stranded fibril). We then use Bayesian inference, where the likelihood function of an aggregate to be x monomers long is given by

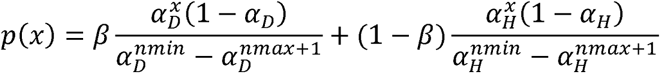

where *x* is the length, *nmin* the minimum aggregate size considered and *nmax* the maximum aggregate size considered. The parameter α is related to the rates of growth, *g*, and removal, *r*, by 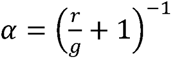

While we use this quantity to fit the aggregate length distribution, in the main text we refer to the relative removal rate *r/g* as that is a more easily interpreted parameter. The minimum size is set to exclude the smallest aggregates where nucleation processes are expected to be important and the assumptions for our length distribution model are thus no longer valid (for details see Cotton et al. (25)). We chose a minimum cutoff, *nmin,* of 500 monomers and a maximum cutoff (*nmax*) of 3000, which prevents outlier effects from single very large aggregates.

To determine α*_H_*, α*_D_* and the fraction of aggregates from diseased cells from experimental data, we perform Bayesian inference, which allows us to directly use each measured length to determine the posterior probability rather than requiring binning and the calculation of histograms. The histograms and “best fit” lines shown in the figures are simply for visualisation.

We assume that, within each extraction method, the healthy and diseased cell states are the same across individuals and stages of the disease (α*_H_* and α*_D_* are global parameters) but allow the fraction of aggregates from diseased cells to differ between individuals. Priors on the parameters are assumed to be constant between 0 and 1. Posteriors are shown in **Supplementary Figure 2**. Even with these restrictive conditions on the parameters, the fits of the experimental data are remarkably good (**see Figure 6b**), showing the distribution from the mean parameters superimposed on a histogram of the data.

## Notes

**Conflict of interest:** Authors declare no conflict of interest.

### Competing Interest Statement

The authors have declared no competing interest.

### Summary of Updates

Some updates on data modelling and figures have been performed.

