## Supplementary figures and images for "Super-resolution microscopy of nanoscopic alpha-synuclein aggregates in brain samples indicates a subset of cells have disrupted protein homeostasis prior to Lewy body formation in Parkinson’s disease"

### Supplemental Figure 1

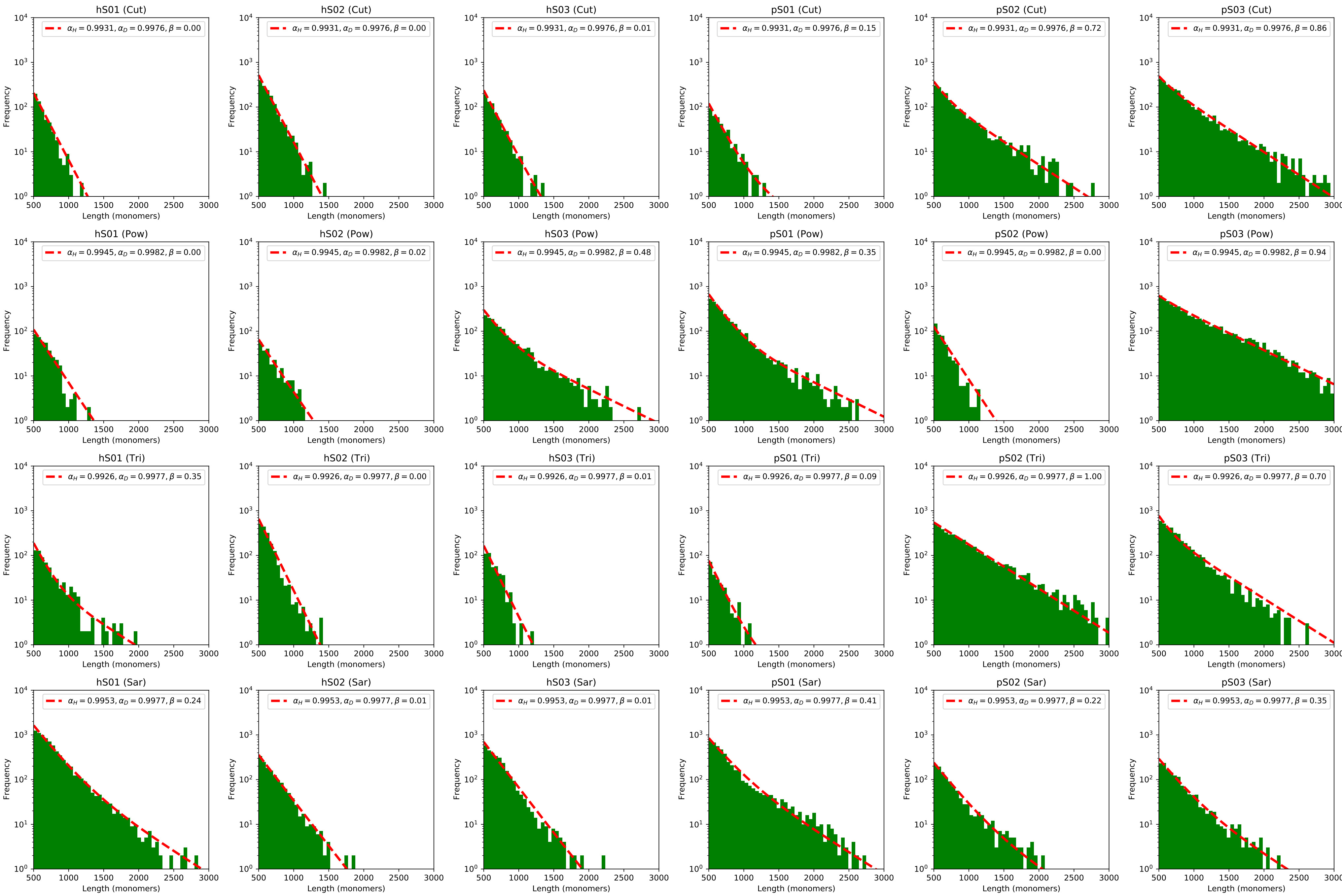

### Supplemental Figure 2

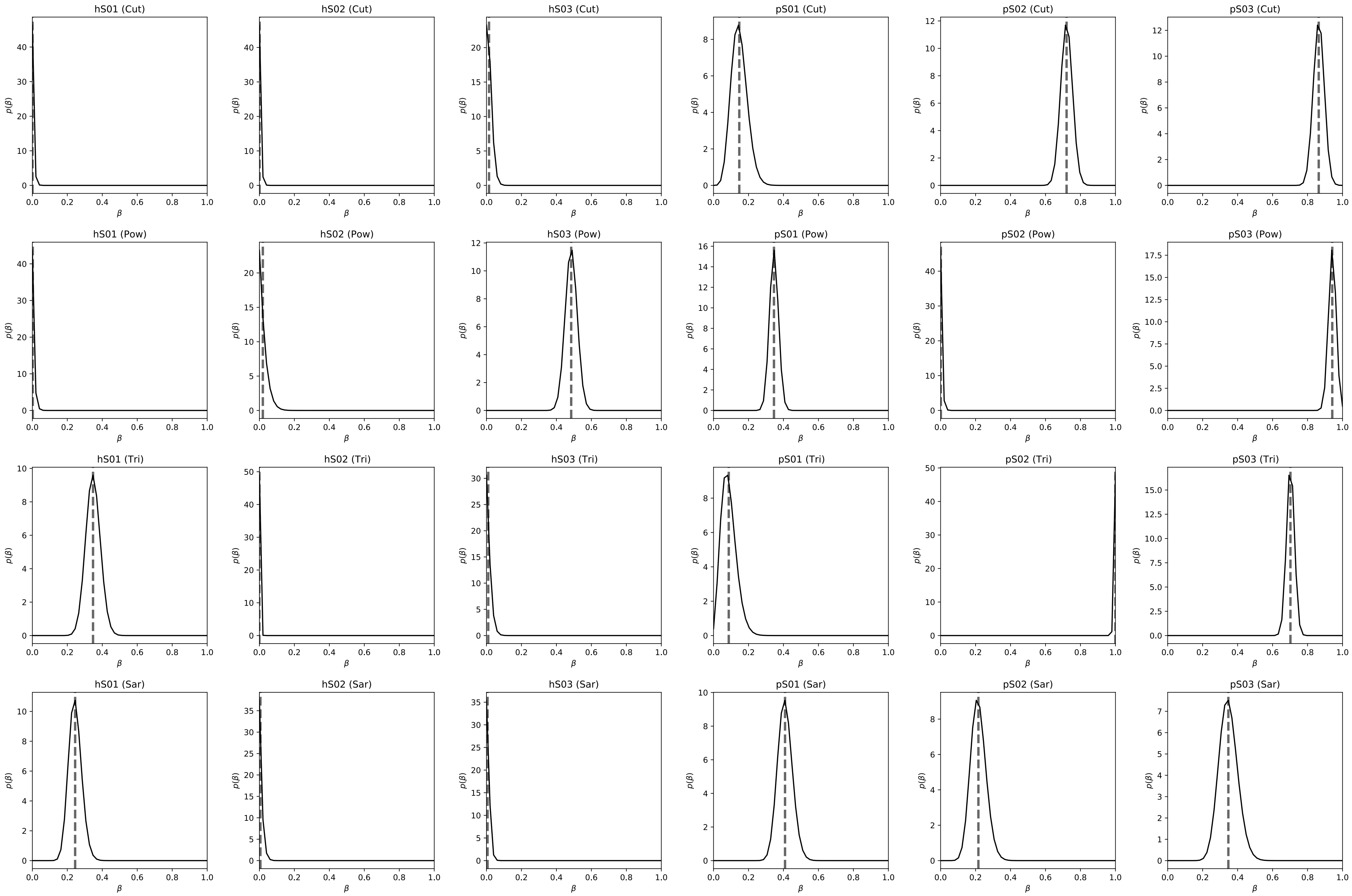

### Supplemental Figure 3

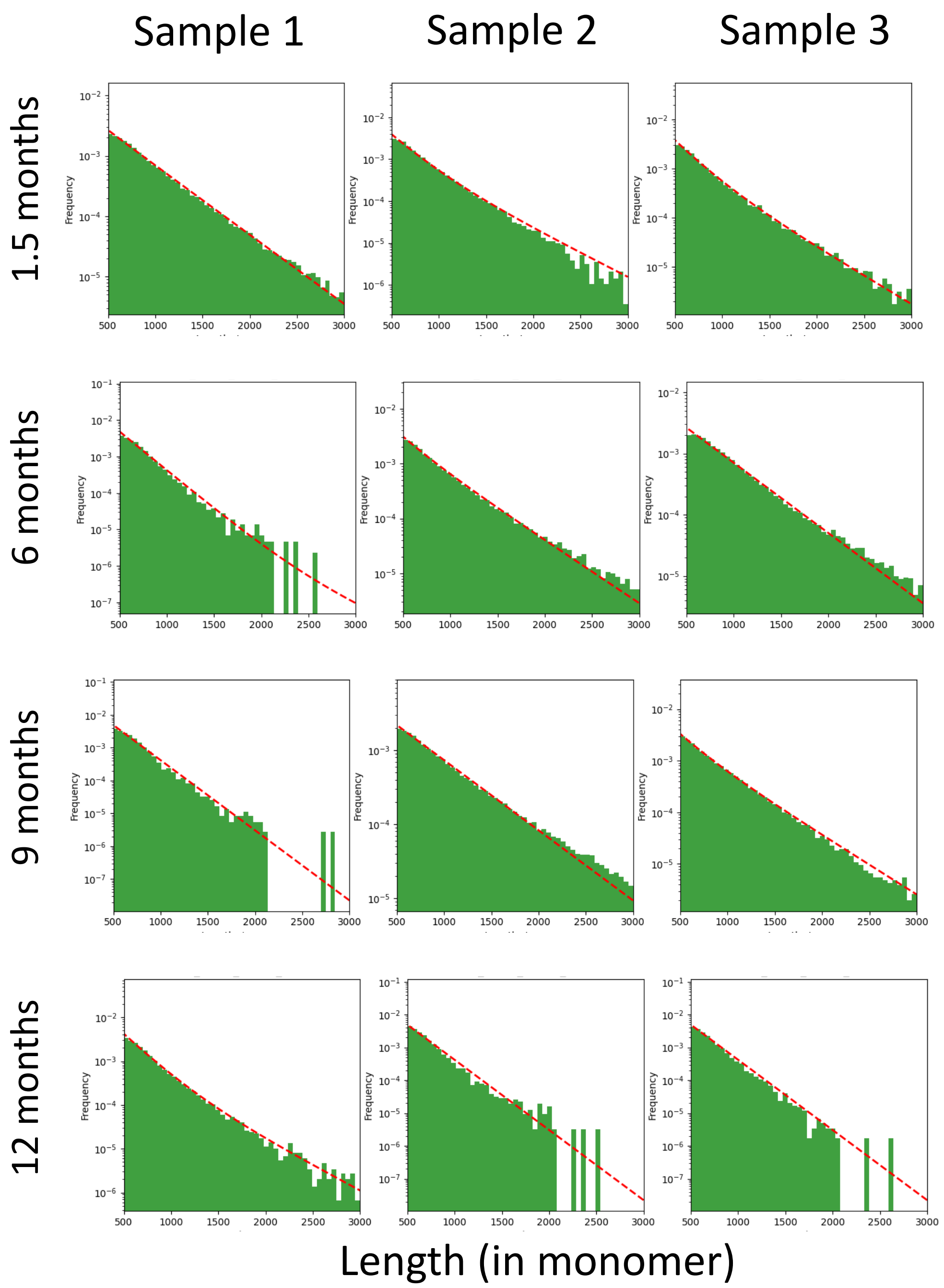

### Supplemental Figure 4

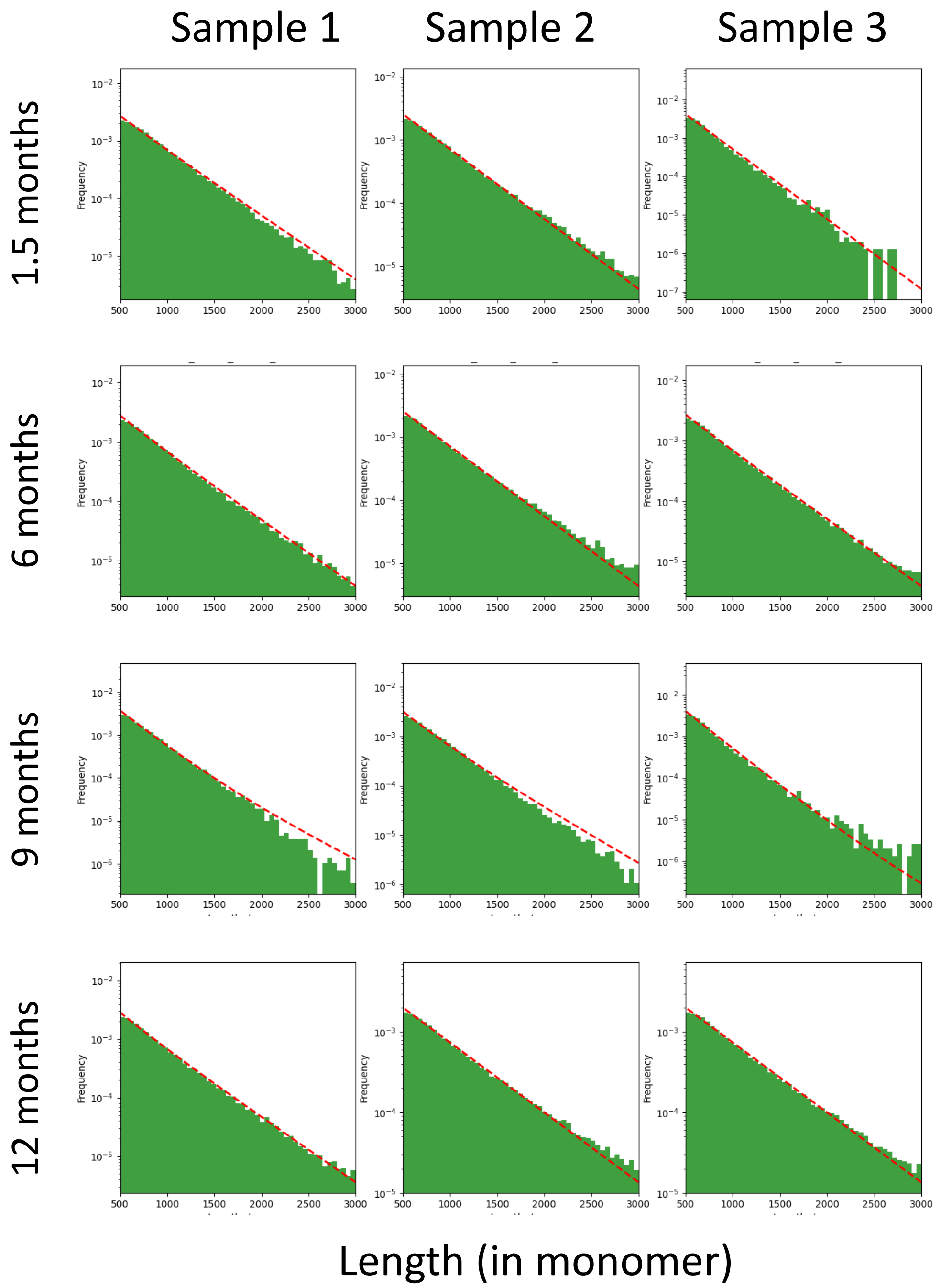

### Supplemental Figure 5

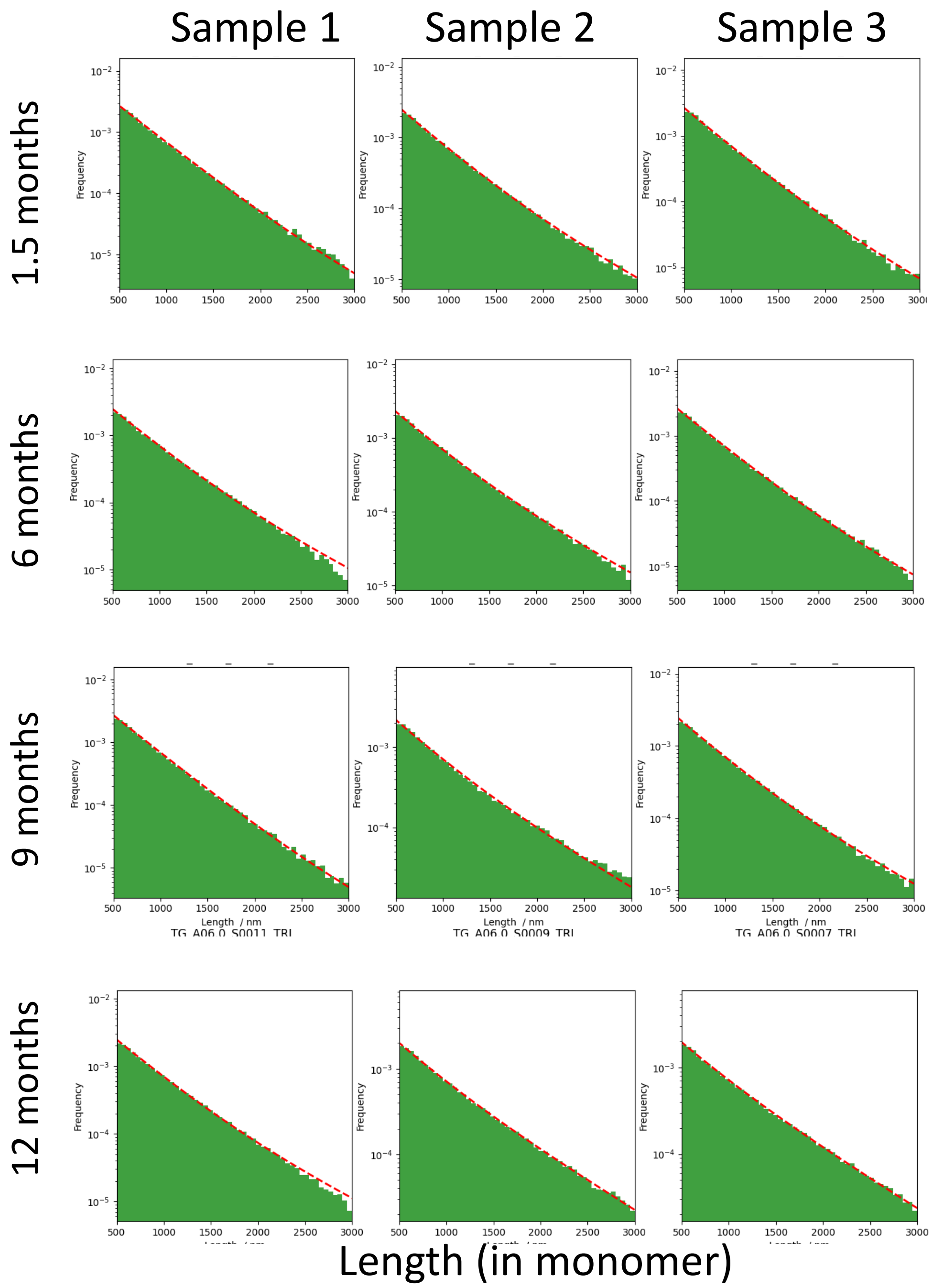

### Supplemental Figure 6

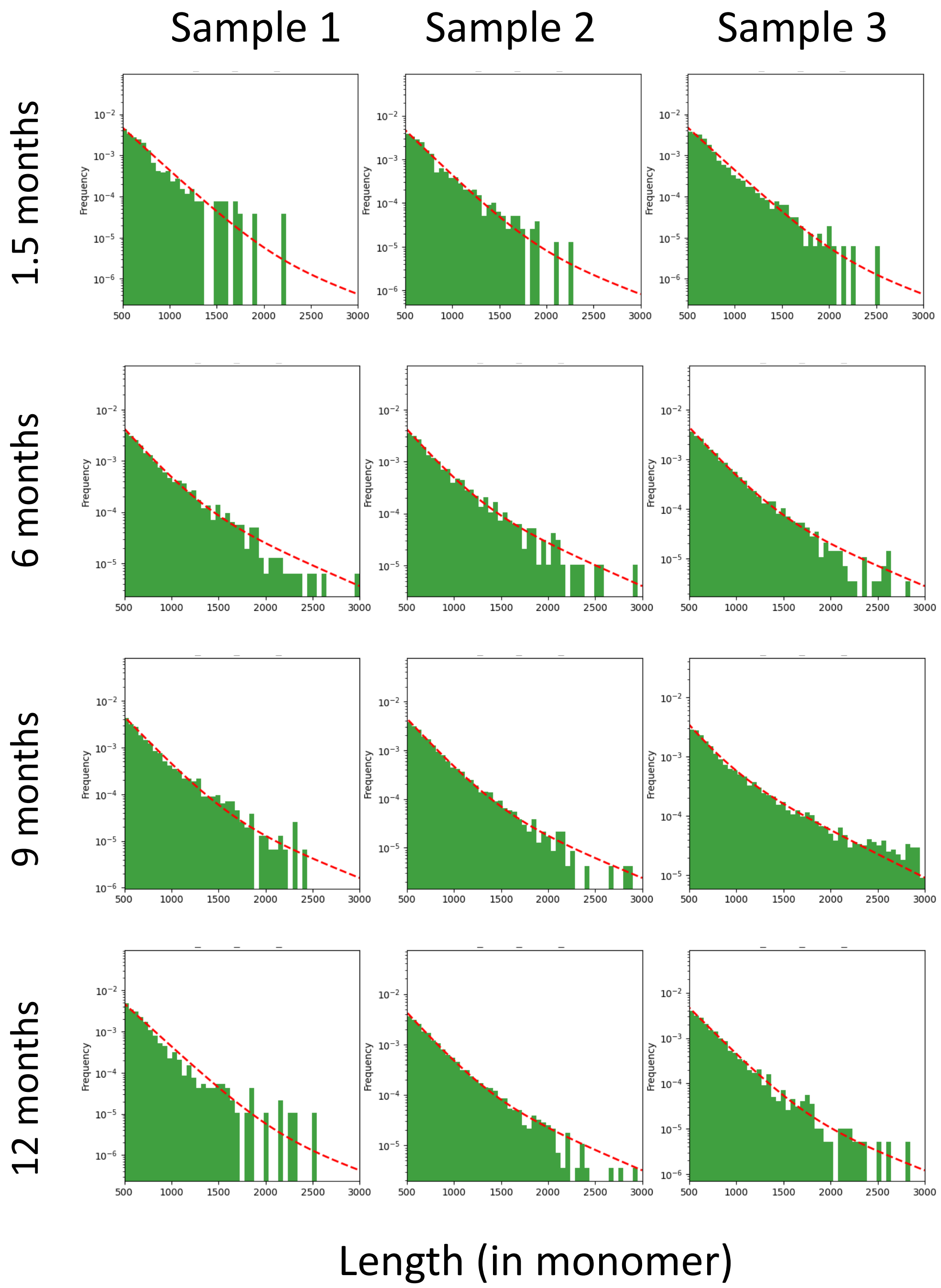
